# The BioRECIPE Knowledge Representation Format

**DOI:** 10.1101/2024.02.12.579694

**Authors:** Emilee Holtzapple, Haomiao Luo, Difei Tang, Gaoxiang Zhou, Niloofar Arazkhani, Casey Hansen, Cheryl A. Telmer, Natasa Miskov-Zivanov

## Abstract

The BioRECIPE (Biological system Representation for Evaluation, Curation, Interoperability, Preserving, and Execution) knowledge representation format was introduced to facilitate seamless human-machine interaction while creating, verifying, evaluating, curating, and expanding executable models of intra- and intercellular signaling. This format allows a human user to easily preview and modify any model component, while it is at the same time readable by machines and can be processed by a suite of model development and analysis tools. The BioRECIPE format is compatible with multiple representation formats, natural language processing tools, modeling tools, and databases that are used by the systems biology community.

**Graphical abstract:** 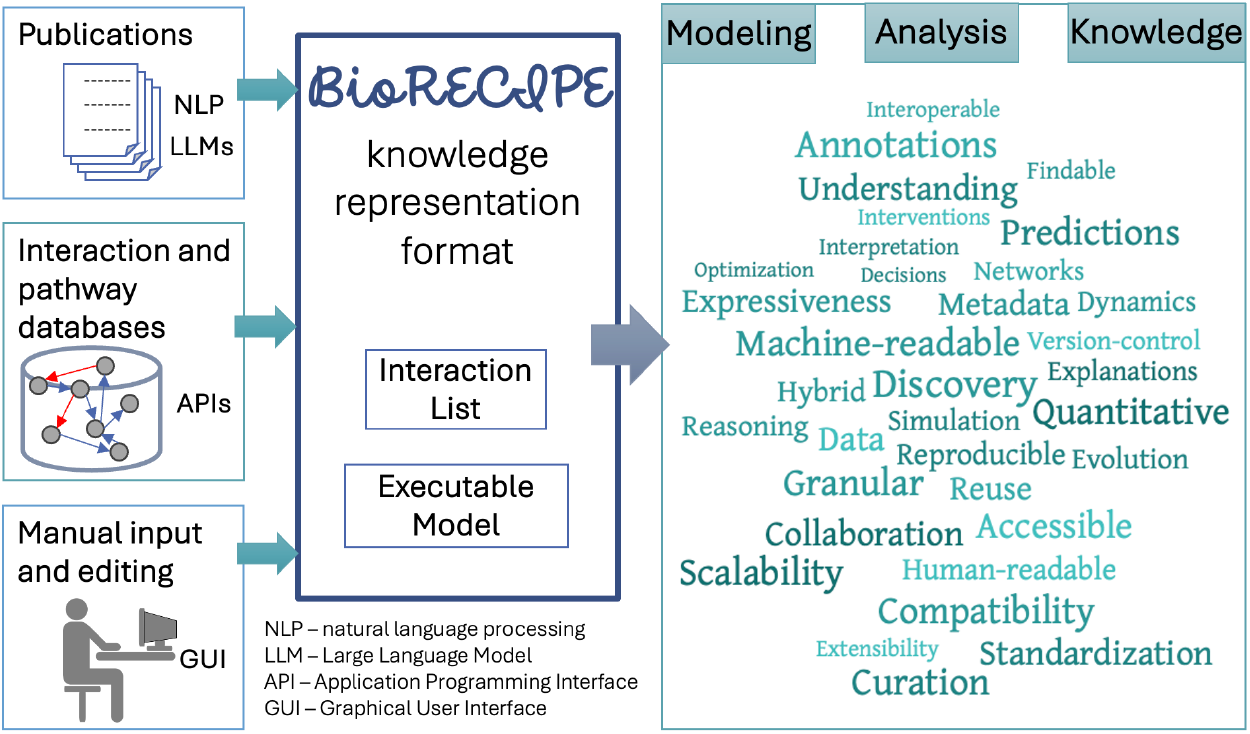

## Introduction

Standard representation formats are essential for creating biological models in a consistent manner. Cell signaling networks (Figure 1A) involve complex events, and as such, there is much variability in the methods used to model these phenomena qualitatively or quantitatively. While formats such as simple-interaction format (SIF), text-based TSV and JSON formats, or graphical formats such as SVG allow for simple representation of cell signaling models, standardized formats can improve accessibility, interoperability, and documentation. These formats help researchers maintain consistency and accuracy in biological modeling.

**Figure 1.**
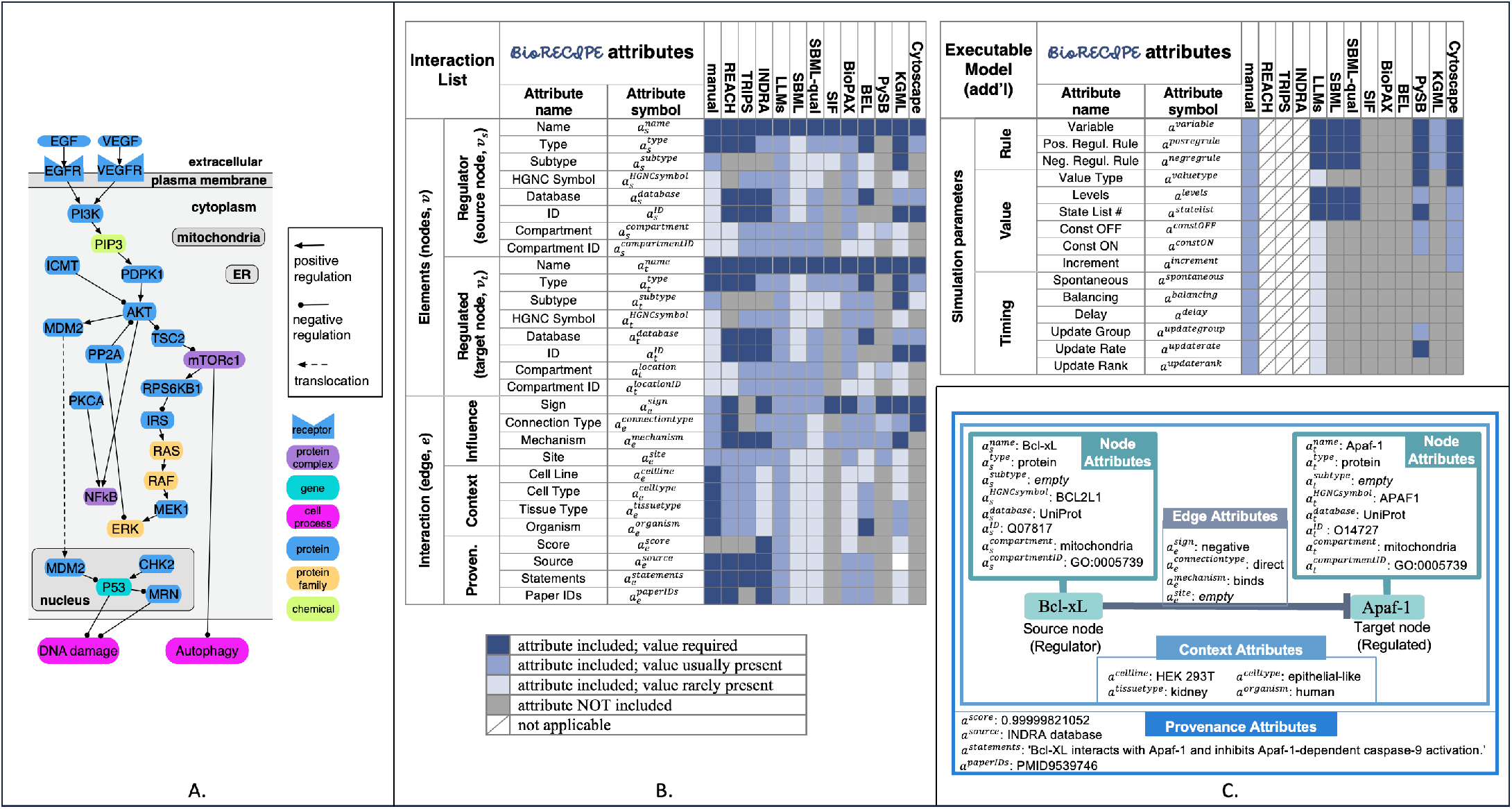
Examples: (A) Pathways, cell compartments, element types, and interactions that can be represented with BioRECIPE; (B) The list of all attributes used by BioRECIPE in Interaction List and Executable Model formats and a summary of whether these attributes are included and required by other formats and tools; (C) An example interaction and interaction attributes that are included in the BioRECIPE’s Interaction List format.

**Figure 2.**
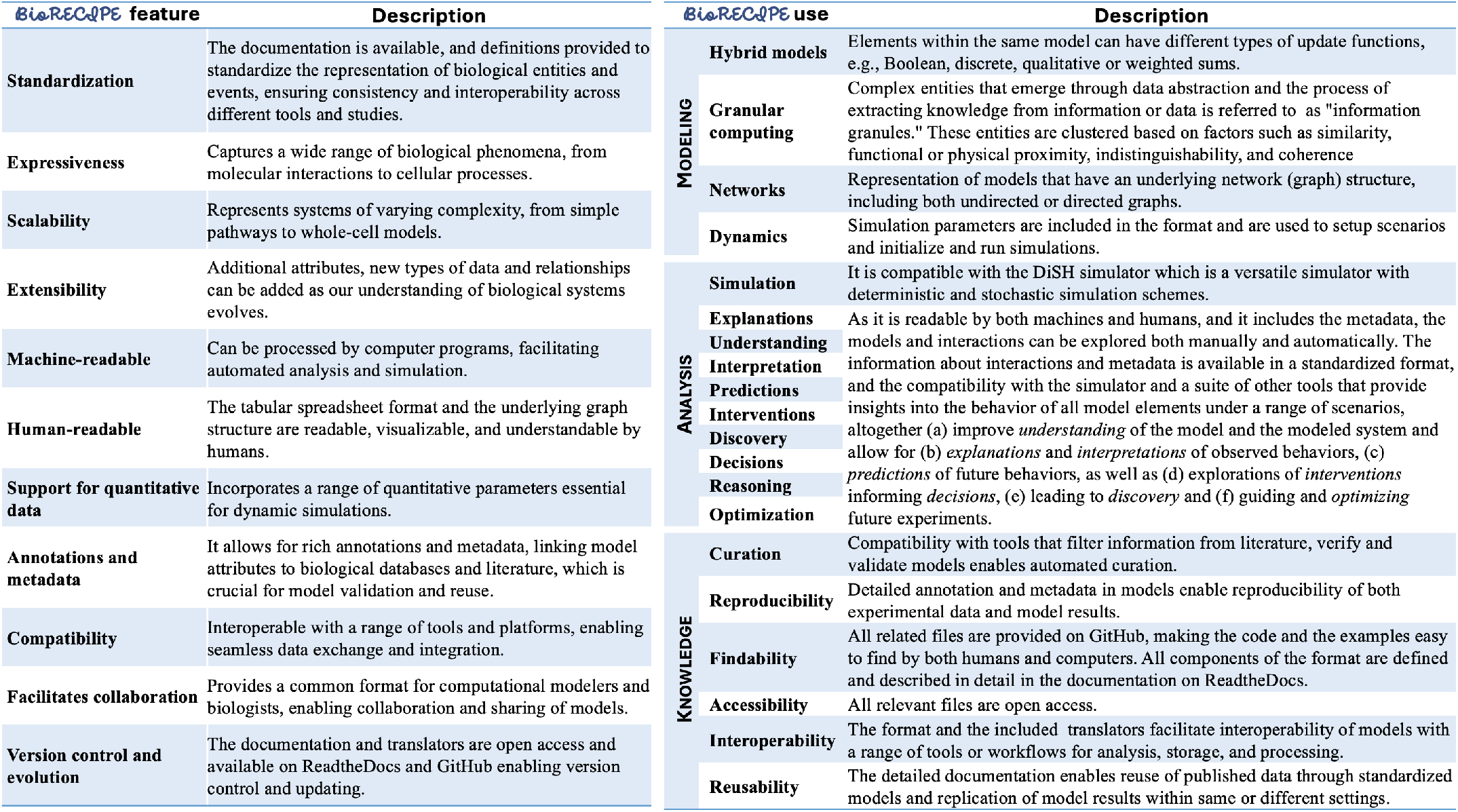
The description of BioRECIPE features and types of models that can be represented with BioRECIPE, model analysis that can be conducted on these models, and the descriptions of how BioRECIPE satisfies the FAIR principles.

One common representation format is the Systems Biology Markup Language (SBML) [1], which seeks to establish a uniform framework for representing models of biological systems. SBML is based on Extensible Markup Language (XML) and is therefore machine-readable. This format uses modules to represent various components within a biological system such as reactions, species, and compartments. SBML supports analysis via ordinary differential equations (ODEs), stochastic simulation algorithm [2], or the reaction rule-based approach (e.g., BioNetGen [3]). SBML format allows for a significant amount of user annotation and provides standardized mechanisms for capturing important information such as starting conditions, context, metadata, and literature sources. However, SBML is not easily interpreted by life scientists without previous exposure to XML. Cell Markup Language (CellML) [4] is also used for standardizing and storing executable models of cellular processes. While it is similar to SBML, the CellML modeling framework requires kinetic parameters for each interaction within the model. Therefore, these two formats differ in scope-CellML is ideal for detailed models of molecular interactions while SBML is more suitable for modeling cell signaling pathways and networks. The Biological Pathway Exchange (BioPAX) [5] provides another standardized format to represent molecular interactions in a signaling pathway. BioPAX supports three levels of representation, with each level offering increased complexity and detail. A simpler representation format, the Biological Expression Language (BEL) [6], represents causal relationships found between biological entities as a triplet statement. Similarly, the INDRA database [7] represents causal relationships between entities in biomedical literature as statements with more detail than BEL. Some interaction and pathway databases may use their own representation format. For example, the KGML representation format [8] was created for storing and standardizing models in the KEGG database.

Model representation formats rely on pre-existing ontologies to standardize individual biological entities and represent biological models. OBO (Open Biological and Biomedical Ontologies) is widely used to represent structured ontologies and controlled vocabularies, including the Gene Ontology (GO) Resource [9]. It is human-readable and allows for the definition of classes, properties, and relationships between terms, making it suitable for standardizing biological models. While not strictly reserved for biological modeling, Web Ontology Language (OWL) is a more powerful and expressive language for creating ontologies. It uses formal logic to create semantic networks and is particularly useful for capturing complex relationships and reasoning in biological models in BioPAX format.

However, there is still a need for a standardized format that may be used for both static and executable models, has translators for seamless conversion into other representation formats, is both human and machine-readable, and can incorporate diverse annotations as well as data. To address this need, here we present the BioRECIPE representation format, which is interoperable with existing interaction, pathway, and model representation formats, and while it utilizes standardized ontologies, it is not dependent on any one ontology (Figure 1B).

## Results

BioRECIPE is a tabular representation format, typically written in a spreadsheet file type. This format can be used by both computational modelers and biology experts to create and modify Interaction List and Executable Model files. It also has a formal structure that can be read, created, updated, and output by computer programs. The detailed BioRECIPE documentation is available as ReadtheDocs pages [10], with instructions to create Interaction Lists and Executable Models in this format.

In BioRECIPE, interactions are represented using the *event-based* Interaction List spreadsheet format, where each biological event is assigned one row, and the columns correspond to attributes of the event participants and the interaction between them. The attributes are selected and organized to allow for detailed curation and an extensible representation of interactions. An example biological interaction, represented as a directed signed edge between two nodes, including node, edge, context, and provenance attributes is illustrated in Figure 1C. Several more examples are included in the Supplement and in the ReadtheDocs documentation.

The BioRECIPE format also provides representation of the static graph structure of models, as well as attributes necessary to study the dynamics, usually through simulations. These executable models are represented in the BioRECIPE format using the *element-based* approach where each element in a model is assigned a row in the model spreadsheet, combining multiple interactions in which the element is the target of the influence. Element update functions are written using a simple notation in the BioRECIPE format, and a number of parameters can be selected to set up simulation scenarios (Figure 1B).

Interactions and models written in the BioRECIPE format can be used with a range of different tools that filter and classify interactions and automatically assemble and analyze models, either by directly using BioRECIPE or by translating models into other formats such as the SBML format (see Supplement). Interactions can be converted to the BioRECIPE format from the output of natural language processing tools or from interaction and pathway databases. Models can be converted from other representation formats and model databases. The ReadtheDocs documentation includes links to example files and translators that convert between BioRECIPE and either other representation formats or inputs and outputs of various tools.

## Conclusions

The BioRECIPE representation format is a valuable tool for systems and synthetic biology that enables comprehensive model curation by both humans and machines. The complexity of cellular signaling pathways and their components necessitates modeling methods that can account for a multitude of details. BioRECIPE allows for all key element and interaction attributes to be included, as well as attributes for simulation, making it compatible with many existing interaction, pathway and model databases, and with a range of tools for extraction of interaction information, model curation, simulation and analysis. This interoperability ensures that researchers can seamlessly integrate BioRECIPE into their existing workflows. Future directions include additional functionality to translate between other existing formats (such as SBOL [11]), or integration of the BioRECIPE representation format as input to model curation and storage platforms (such as CellCollective [12] or NDEx [13]). These platforms play a crucial role in managing and disseminating computational models, and the integration with BioRECIPE could further streamline the process of model sharing and collaboration within the scientific community.

