## Supplemental Materials for "The BioRECIPE Knowledge Representation Format"

### Supplementary Material

| | | BioRECIPE attributes: Attribute name | Attribute value examples: Interaction 1 | Interaction 2 | Interaction 3 | Interaction 4 | Interaction 5 |
|---|---|---|---|---|---|---|---|
| Elements (nodes, $v$) | Regulator (source node, $v_s$) | Name | CK1 | mTOR | Akt | Resveratrol | RAS |
| | | Type | protein | protein | protein family | chemical | protein family |
| | | Subtype | kinase | kinase | kinase | antioxidant | GTPase |
| | | HGNC Symbol | CSNK1A1 | MTOR | AKT1,AKT2, AKT3 | N/A | HRAS, KRAS, NRAS |
| | | Database | UniProt | UniProt | FamPlex | CHEBI | PFAM |
| | | ID | P48729 | P42345 | AKT | 445154 | PF00071 |
| | | Compartment | cytoplasm | cytoplasm, nucleus | cytoplasm, membrane | N/A | other |
| | | Compartment ID | GO: 0005737 | GO:0005737, GO:0005634 | GO:0005737, GO:0016020 | N/A | GO:0016020 |
| | Regulated (target node, $v_t$) | Name | APC | Chk1 | GSK3beta | PTEN | p110gamma |
| | | Type | protein | protein | protein | protein | protein |
| | | Subtype | tumor suppressor | kinase | kinase | phosphatase | enzyme |
| | | HGNC Symbol | APC | CHEK1 | GSK3B | PTEN | PIK3CG |
| | | Database | UniProt | UniProt | UniProt | UniProt | UniProt |
| | | ID | P25054 | O14757 | P49841 | P60484 | P48736 |
| | | Compartment | cytoplasm | cytoplasm, centrosome | cytoplasm, nucleus, membrane | cytoplasm, nucleus | other |
| | | Compartment ID | GO: 0005737 | GO: 0005737, GO:0005813 | GO:0005737, GO:0005634, GO:0016020 | GO:0005737, GO:0005634 | GO:0016020 |
| Interaction (edge, $e$) | Influence | Sign | positive | positive | negative | positive | positive |
| | | Connection Type | direct | indirect | indirect | indirect | indirect |
| | | Mechanism | phosphorylation | amount | phosphorylation | transcription | activation |
| | | Site | S1504, S150, S1507, S1510 | N/A | S9 | N/A | N/A |
| | Context | Cell Line | SW480 | cellosaurus:CVCL_0045 | CHO | LNCaP, DU145 | COS-7 |
| | | Cell Type | colorectal cancer | embryonic | epithelial | prostate cancer | fibroblast-like |
| | | Tissue Type | large intestine | kidney | ovarian | brain, lymph node | kidney |
| | | Organism | human | human | human, mouse, hamster | human | human, monkey |
| | Proven. | Score | INDRA: 0.999 | STRING: 0.763, INDRA:0.982 | STRING: 0.996, INDRA:0.999 | STITCH: 0.963, INDRA:0.999 | INDRA:0.997 |
| | | Source | literature, expert, database | literature, expert, database | literature, expert, database | literature, expert, database | literature, expert, database |
| | | Statements | (1)* | (2)* | (3)* | (4)* | (5)* |
| | | Paper IDs | PMC2654145, PMID11487578 | PMC4381605 | PMC1403772, PMC3535741 | PMC3181262, PMC2957324 | PMC2652403 |

A.

| | Standard, Database, Tool | Format | To BioRECIPE | From BioRECIPE | Translator Description |
|---|---|---|---|---|---|
| Standard representation formats | SBML | RDF/XML | ✓ | ✓ | Translation to BioRECIPE *Executable Model* and from BioRECIPE *Interaction List* |
| | SBML-qual | RDF/XML | ✓ | ✓ | Translation to and from BioRECIPE *Executable Model* |
| | SIF | TXT | ✓ | ✓ | Translation to and from BioRECIPE *Interaction List* and from BioRECIPE *Executable Model* |
| | BioPAX | RDF/OWL, SBML | ✓ | ✓ | Conversion from and to BioPAX files can be done through SBML translation to and from BioRECIPE |
| | BEL | TXT, (INDRA) | ✓ | ✓ | Conversion from and to BEL statements through INDRA statements |
| | PySB | SBML | ✓ | ✓ | Translation from and to PySB files can be done through the SBML translation to and from BioRECIPE |
| Databases | KEGG | KGML, SBML | ✓ | ✓ | Conversion from and to KGML files through the SBML translation to and from BioRECIPE |
| | REACTOME | SBML, BioPAX | ✓ | ✓ | See SBML and BioPAX conversion |
| | Pathway Commons | SIF, BioPAX | ✓ | ✓ | See SIF and BioPAX conversion |
| | NDEx | SIF, BEL (INDRA), BioPAX | ✓ | ✓ | See SIF, BEL, and BioPAX conversion |
| | BioModels | SBML, SBML-qual | ✓ | ✓ | See SBML and SBML-qual conversion |
| External tools (and databases) | Cytoscape | SIF, CX (INDRA) | ✓ | ✓ | See SIF conversion or conversion through INDRA statements |
| | Cell Collective | SBML-qual | ✓ | ✓ | See SBML-qual conversion |
| | CellNetAnalyzer | SBML | ✓ | ✓ | See SBML conversion |
| | CellDesigner | SBML | ✓ | ✓ | See SBML conversion |
| | INDRA | JSON | ✓ | ✓ | Conversion to and from BioRECIPE *Interaction List* |
| | REACH | JSON | ✓ | N/A | Conversion to BioRECIPE *Interaction List* |
| | TRIPS | XML | ✓ | N/A | Conversion to BioRECIPE *Interaction List* |
| Internal tools | DiSH | BioRECIPE | ✓ | ✓ | Uses BioRECIPE format at input |
| | FLUTE | BioRECIPE | ✓ | ✓ | Uses BioRECIPE format at input |
| | VIOLIN | BioRECIPE | ✓ | ✓ | Uses BioRECIPE format at input |
| | CLARINET | BioRECIPE | ✓ | ✓ | Uses BioRECIPE format at input |
| | ACCORDION | BioRECIPE | ✓ | ✓ | Uses BioRECIPE format at input |
| | Model-List converter | BioRECIPE | ✓ | ✓ | Converts between Interaction List and Executable Model formats |

B.

Figure S1.

A. Example attribute values extracted from five sentences. Sentences used:
*(1) “This may be analogous to parallel mechanisms that promote GSK3 phosphorylation of beta-catenin in the absence of Wnt stimulation, such as by GSK3 and CK1 phosphorylation of Axin and APC.”
(2) “mTOR inhibition in HEK293 cells significantly reduced the total Chk1 level"
(3) “Therefore, given that Akt phosphorylates and inactivates GSK3beta, we hypothesized that Akt dependent inactivation of GSK3beta might be responsible for Notch potentiation.”
(4) “Our results demonstrate that resveratrol induced the expression of PTEN…”
(5) “Ras activates p110gamma at the level of the membrane, by allosteric modulation and/or reorientation of the p110gamma…”

B. Formats for which we have created translators and converters, and databases and tools with which BioRECIPE is either directly compatible or can be converted to and from the input/output formats for these databases and tools (all the links to translators are available in ReadtheDocs documentation for BioRECIPE).
